# Transspecies beak color polymorphism in the Darwin’s finch radiation

**DOI:** 10.1101/2021.01.19.426595

**Authors:** Erik D. Enbody, C. Grace Sprehn, Arhat Abzhanov, Huijuan Bi, Mariya P. Dobreva, Owen G. Osborne, Carl-Johan Rubin, Peter R. Grant, B. Rosemary Grant, Leif Andersson

**Affiliations:** Department of Medical Biochemistry and Microbiology, Uppsala University, SE-751 23 Uppsala, Sweden; Department of Life Sciences, Imperial College London, Silwood Park Campus, SL5 7PY Ascot, UK; School of Natural Sciences, Bangor University, Environment Centre Wales, Deiniol Road, Bangor, LL57 2UW, UK; Department of Ecology and Evolutionary Biology, Princeton University, Princeton, New Jersey 08544, USA; Department of Animal Breeding and Genetics, Swedish University of Agricultural Sciences, SE-75007 Uppsala, Sweden; Department of Veterinary Integrative Biosciences, Texas A&M University, College Station, Texas 77843-4458, USA

## Abstract

Carotenoid-based polymorphisms are widespread in populations of birds, fish, and reptiles^1^, but little is known of how they affect fitness and are maintained as species multiply^2^. We report a combined field and molecular-genetic investigation of a nestling beak color polymorphism in Darwin’s finches. Beaks are pink or yellow, and yellow is recessive^3^. Here we show that the polymorphism arose in the Galápagos approximately half a million years ago through a regulatory mutation in the *BCO2* gene, and is shared by 14 descendant species. The frequency of the yellow genotype is associated with cactus flower abundance in cactus finches, and is altered by introgressive hybridization. The polymorphism is most likely a balanced polymorphism, maintained by ecological selection pressures associated with diet, and augmented by occasional interspecific introgression. Polymorphisms that are hidden as adults, as here, may contribute to evolutionary diversification in underappreciated ways in other systems.

## Main Text

Adaptive radiations are groups of related organisms that have diversified relatively rapidly from a common ancestor^4,5^. A striking feature of some radiations is that polymorphic variation within species is shared among related species (reviewed in^2^). The question this raises is how the same polymorphisms persist among related species when the rarer morph may be lost in the speciation process due to drift^2,6–8^. Their origin can potentially be accounted for by shared ancestral variation, repeated mutation and/or introgression, and their maintenance can be explained by negative frequency-dependent selection, heterozygous advantage or spatiotemporal fluctuations in selective pressures^2,7,8^. Distinguishing between these alternatives requires an understanding of the genetic basis for phenotypic variation, phylogenetic history, and fitness variation in natural populations. This has been rarely accomplished, in part because in many cases polymorphisms are associated with large genomic regions containing many genes of uncertain functional importance (e.g.^9^). Here we report a unique form of shared polymorphism in the radiation of Darwin’s finches (Thraupidae) on the Galápagos and Cocos islands. We identify its genetic basis and phylogenetic origin, and take advantage of uniquely banded individuals on one island to quantify relative fitness in order to address the question of long-term maintenance of the two morphs.

A beak color polymorphism in nestlings has been documented in eight species of Darwin’s finches (one *Camarhynchus* and seven *Geospiza^10^).* The beaks of nestlings are either pink or yellow (Fig. 1A). Pedigrees on Daphne Major island^3^ and Genovesa^11^ show that the yellow phenotype is recessively inherited, but the causal gene is unknown. Increasing melanin deposition in the beak obscures phenotypic expression several weeks after fledging (the phenotype is recognizable in some juveniles for up to three weeks^10^) and the beak is fully melanized in breeding birds. Frequencies of the morphs in two species of ground finches on Genovesa and two other species on Daphne Major remained stable for more than a decade^3^.

**Fig. 1:**
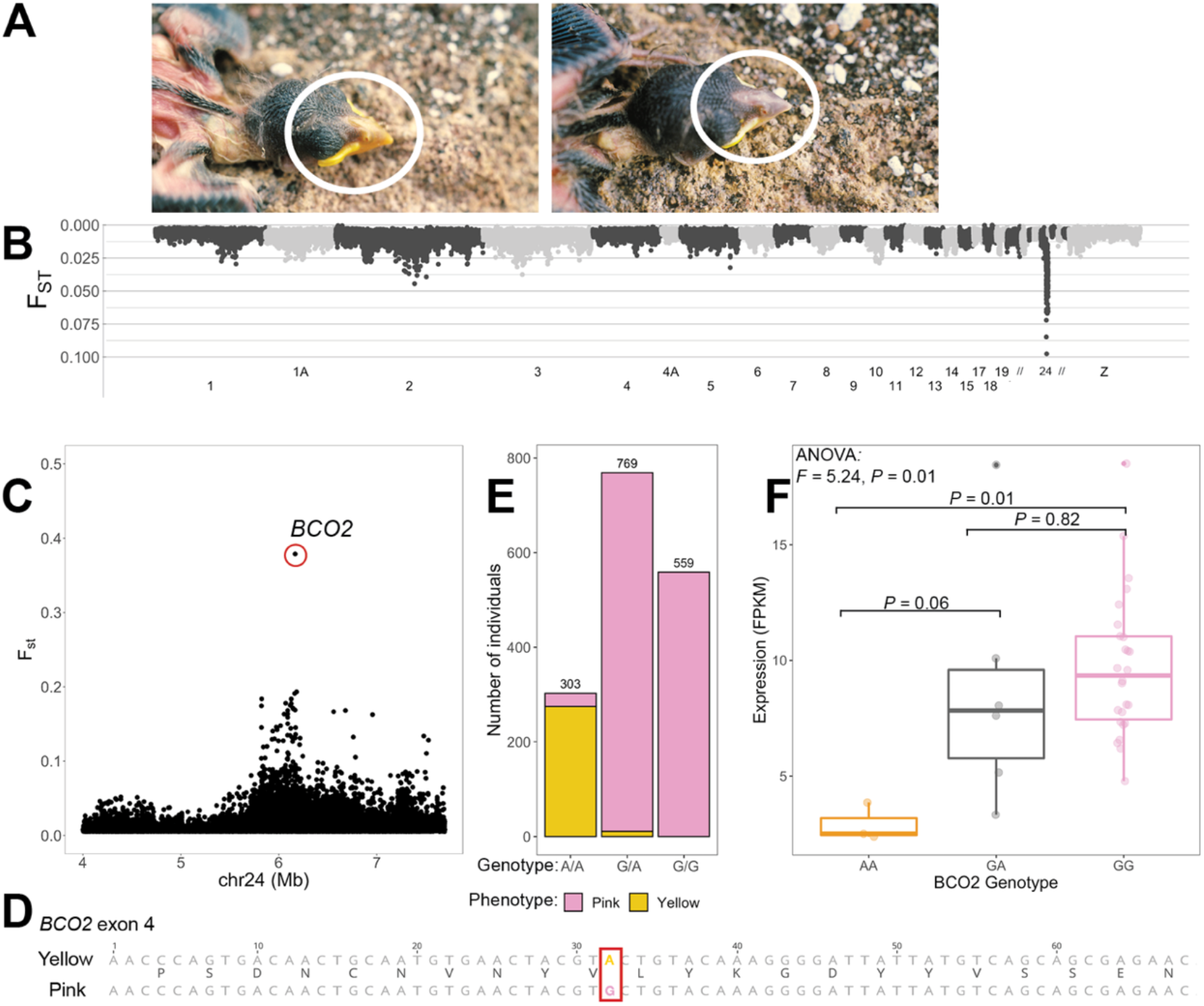
Genetic basis for a beak polymorphism in Darwin’s finches. **(A)** *G. magnirostris* nestlings with the yellow beak phenotype (left) or pink beak phenotype (right). Images courtesy of P.R. Grant. **(B)** Whole-genome F_ST_ scan using 10kb, non-overlapping, windows contrasting 97 pink with 97 yellow individuals. **(C)** Per-site F_ST_ estimates in the region showing strong differentiation on chromosome 24 and a single SNP (F_ST_ = 0.38) within exon 4 of *BCO2* is highlighted. **(D)** Alignment of the first 68bp in exon 4 of *BCO2* between a yellow and pink beak individual, with translated amino acids indicated between alignments. The high F_ST_ SNP from **(C)** is highlighted. **(E)** Phenotype to genotype matching using 1,631 individuals of known phenotype. Sample sizes for each group are marked above bars. **(F)** Fragments Per Kilobase per Million mapped reads (FPKM) of *BCO2* in the upper beak of ground (n = 3 species), tree (n = 2 species), and vegetarian finch embryos, contrasted across known genotypes. *P*-values for Tukey’s post-hoc analysis is noted above each individual comparison and boxplot hinges correspond to the first and third quantiles, center line is median, and whiskers mark 1.5x the interquartile range. Raw data are shown as points and jittered.

We used samples collected as part of long-term monitoring of three species known to carry the polymorphism on Daphne Major: *Geospiza fortis, G. scandens,* and *G. fuliginosa.* We sequenced whole genomes at low coverage of individuals of known phenotype as nestlings (n = 213 for each color morph; mean depth = 2X, Extended Data Fig. 1) to search for the genetic basis of the polymorphism. We generated genotype likelihoods using ANGSD^12^ and used window-based F_ST_ outlier analysis (comparing beak phenotypes, n = 97 each, of *G. fortis* samples) to search for genomic regions associated with beak color. We discovered a small region on chromosome 24 harboring extremely deviant F_ST_ values in a region overlapping the carotenoid-cleaving beta-carotene oxygenase 2 gene (*BCO2*, Fig. 1B). Mutations downregulating BCO2 activity result in increased deposition of carotenoids and pigmented phenotypes in birds, mammals, and reptiles^13-15^. Close inspection of this region in all 416 finches identified a single exonic single nucleotide polymorphism (SNP) with F_ST_ = 0.38, nearly the maximum value expected for a recessive marker (Fig. 1C). This SNP leads to a synonymous change 32bp into exon 4 of *BCO2*. We used high coverage sequencing data for 16 pink and 8 yellow individuals to search for linked SNPs or structural variants in the vicinity but found none and confirmed the phenotype association with the exon 4 SNP (Extended Data Fig. 2).

In order to confirm the association for our SNP of interest we designed a TaqMan SNP assay to genotype additional individuals. In total, we genotyped 2,890 individuals captured on seven Galápagos islands and Cocos Island from nine species (including hybrids) during the period 1975 to 2011. Our dataset consisted of 1,631 individuals of known phenotype (predominantly *G. fortis* and *G. scandens,* Extended Data Table 1), the observed genotype matched the predicted genotype based on phenotype for 98% of them (Fig. 1E). Mismatched pink phenotype with yellow genotype could be the result of mis-phenotyping or limited nutrition^16^, but mismatched individuals were notably often clustered in families (Extended Data Fig. 3), suggesting a possible genetic contribution. No homozygous pink genotypes showed the yellow phenotype.

The functional importance of the observed synonymous change is uncertain, but codon usage bias can be under strong selection^17^ and have functional consequences, including altering transcription^18^. We found that yellow homozygotes showed significantly lower *BCO2* expression compared to pink homozygotes in the upper beak of developing embryos (Fig. 1F). We used the transcription factor affinity prediction (sTRAP^19^) web tool to evaluate the hypothesis that transcription factor binding sites are affected by the identified SNP. The pink allele has a significantly greater predicted affinity for nine transcription factors compared to the yellow allele, while none were predicted to bind better to the yellow over the pink allele (Extended Data Table 2). This is consistent with the observed difference in transcriptional regulation (Fig. 1F). The greatest difference was predicted for hypoxia-inducible factor 1 (HIF1, *P*_pink_ = 0.0086 vs. *P*_yellow_ = 0.27). RT-PCR analysis revealed no variation in splice forms when amplifying across exons in alternative homozygous individuals (Materials and Methods).

Our phylogenetic analysis parsimoniously shows that it is unlikely that the yellow allele was brought to the Galápagos by the founders of the radiation, but instead first appeared in the Galápagos approximately half a million years ago^20^ and that the ancestral state is homozygous pink (Fig. 2). The yellow allele is not present in *G. septentrionalis,* despite the fact that all individuals of this species have yellow beaks, as well as yellow legs and yellow skin, that are maintained into adulthood^10^. This unique phenotype is likely associated with another mutation in *BCO2* or other gene affecting carotenoid metabolism that makes the synonymous change redundant.

**Fig. 2:**
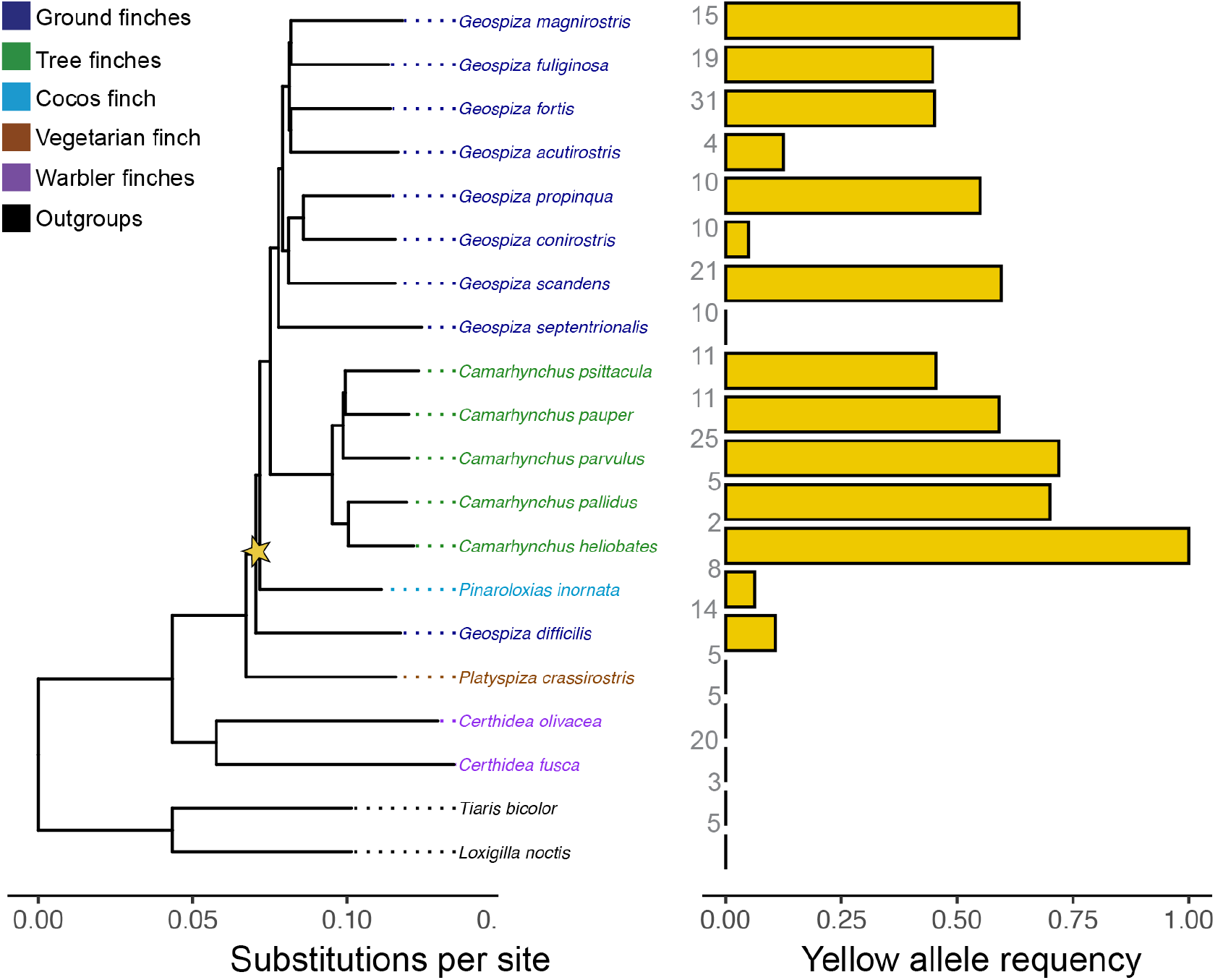
Frequency of the yellow allele across the Darwin’s finch radiation. Left, a species tree for Darwin’s finches (reproduced from^21^), colored by genera, with the parsimonious origin of the yellow allele marked with a yellow star. Right, the frequency of the yellow allele (A) in all finch species with the number of individuals genotyped marked along the vertical axis (based on whole genome sequencing data from^20,21^.

Nucleotide polymorphisms that are shared among 14 or more species of Darwin’s finches, like the *BCO2* polymorphism, make up roughly 5% of all polymorphic sites. Thus, the *BCO2* polymorphism lies in the tail of the distribution of polymorphic sites that show extensive transspecies polymorphism in the phylogeny. Such long-term persistence of a polymorphism, both among (Fig. 2) and within (Fig. 3) species, implies some form of balancing selection (reviewed in^8^). On the small island of Daphne Major, the cactus-feeding specialist, *G. scandens* (Fig. 3A), had a higher yellow genotype frequency (~30%) than *G. fortis* (~20%, Fig. 3C) for many years, matching previous estimates deduced from phenotypes^3^. However, the two homozygous genotypes display strong annual variation (coefficient of variation 33.7 ± 10.4 for the yellow genotypes and 28.6 ± 8.5 for the pink homozygotes, n = 26 years), resulting from a shift in yellow genotype frequencies beginning at the turn of the century (Fig. 3C). The trends in alternate homozygotes are in opposite directions (adjusted R^2^ value = 0.51, *P* <0.0001), and are so strong that the relative frequencies of the homozygotes became reversed (Extended Data Fig. 6). Variation helps to identify possible causal factors by correlation. The main factor affecting homozygote frequencies is introgression of alleles from *G. fortis.* Introgression introduces the pink allele at a higher frequency than the yellow allele (Fig. 3C), thereby increasing the chances of homozygous pink genotypes being formed and decreasing the chances of yellow genotypes being formed. Consistent with this reasoning, annual variation in proportions of microsatellite-identified hybrids^22^ is negatively related to the frequency of yellow genotypes (adj R^2^ = 0.51, *P*<0.0001). Hybridization is bidirectional but unequal: *G. fortis* donates more than it receives (Supplemental text).

**Fig. 3:**
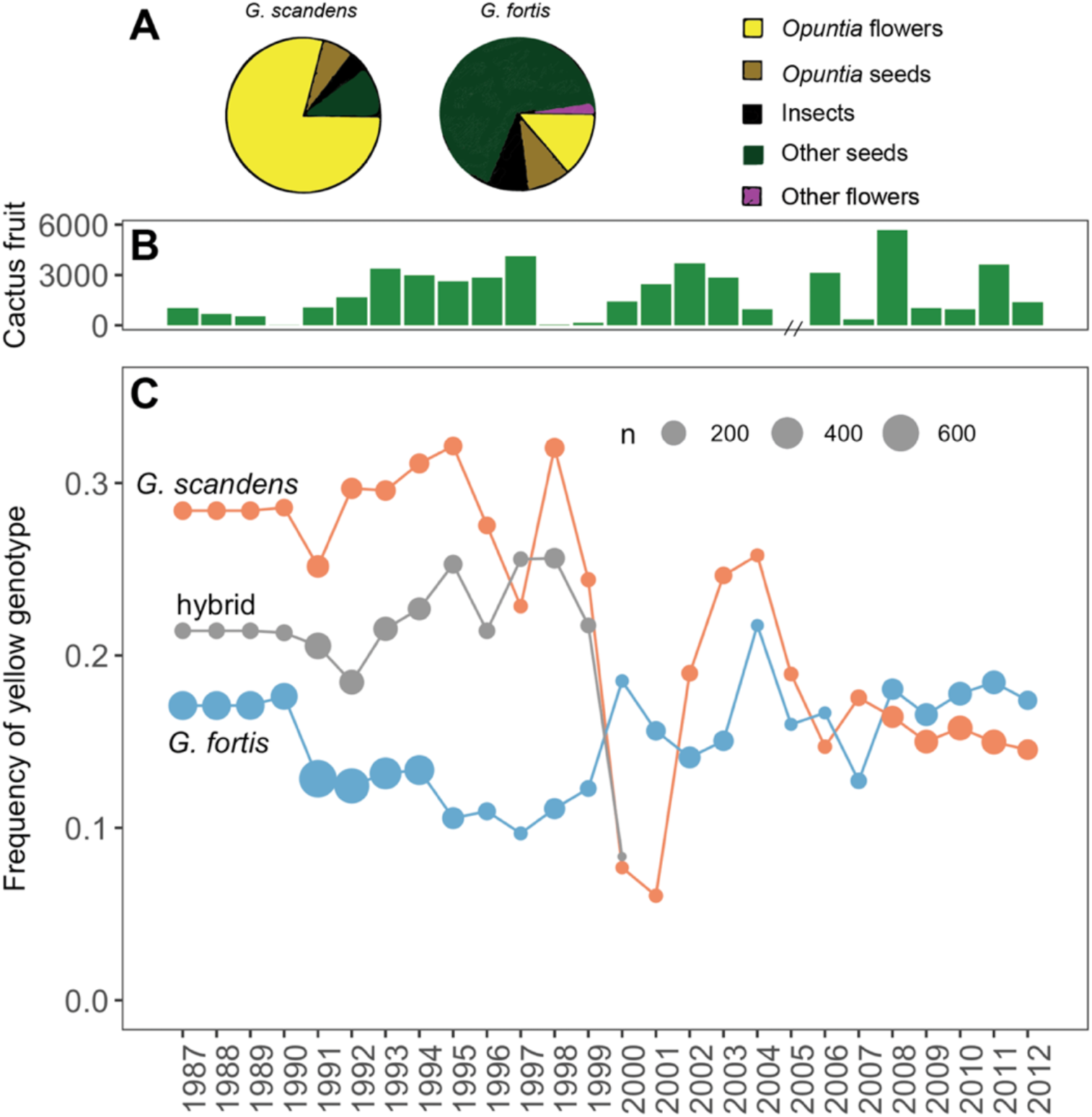
Changes in yellow genotype frequency over time in relation to cactus abundance in two Darwin’s finch species and their hybrids on Daphne Major. **(A)** Proportion of diet during early breeding in *G. scandens* and *G. fortis* (reproduced from^23^). **(B)** Annual maximum *Opuntia* cactus fruit availability on Daphne Major. No data were collected in 2005. **(C)** Frequency of the yellow (A/A) genotype over 26 years for *G. fortis* (blue) and *G. scandens* (orange) and hybrids (F_1_ and backcrosses) in grey, points are scaled by sample size.

The *BCO2* polymorphism might affect fitness in several ways, in particular since *BCO2* has an essential role in Vitamin A metabolism^24^. Nestling color variation might have a signaling role if nestlings are fed preferentially by their parents^25^. Observations made during half hour nest watches, and parental feeding of juveniles out of the nest, gave no indication of preferential feeding^10^. Another possibility is that carotenoid deposition early or late in life could improve immune function^26^ or lead to a reduction in oxidative stress^27^. However, we did not find evidence for an improved immune defense among yellow homozygotes in relation to an outbreak of the pox virus in 2008 (^28^, Extended Data Fig. 8, χ^2^ = 0.98, *P* = 0.61, df = 2). A third possibility is that deposition in beaks avoids a toxic accumulation of metabolic breakdown products of circulating carotenoids^24^. Deposited and stored carotenoids can then be metabolized later at a time of lower intake^29^. It is known that excess carotenoid accumulation impairs muscle function in other bird species^30^.

All species of Darwin’s finches obtain carotenoids by feeding on pollen and/or herbivorous insects, mainly Lepidoptera larvae. In particular, cactus pollen accounts for a large fraction of the breeding season diet of the cactus finches, *G. scandens* and *G. propinqua*^3,11^; Fig. 3A). On Daphne Major, a comparison of the two years with the largest samples of nestlings illustrates the association between yellow allele fitness in *G. scandens* and cactus supply. Individuals with the homozygous yellow genotype had high rates of survival in 1991, a year of abundant flower and fruit production, compared to 1998 when cactus products were almost absent (Fig. 4A). Thus, according to this contrast, the yellow homozygous genotype in *G. scandens* may have a survival advantage in years of high cactus availability (also observed in years with only phenotype data, Extended Data Fig. 7). The unusually low supply of cactus flowers and fruits coupled with high mortality of yellow homozygous individuals continued into 1999 (Fig. 3). Possibly, yellow genotype individuals experienced a disadvantage following from the interruption to the essential Vitamin A metabolism mediated by downregulation of *BCO2* expression. Low survival was followed by hybridization with *G. fortis*, with the result that the frequencies of the yellow genotype in *G. scandens* and *G. fortis* converged (Fig. 3C, Extended Data Fig. 5).

**Fig. 4:**
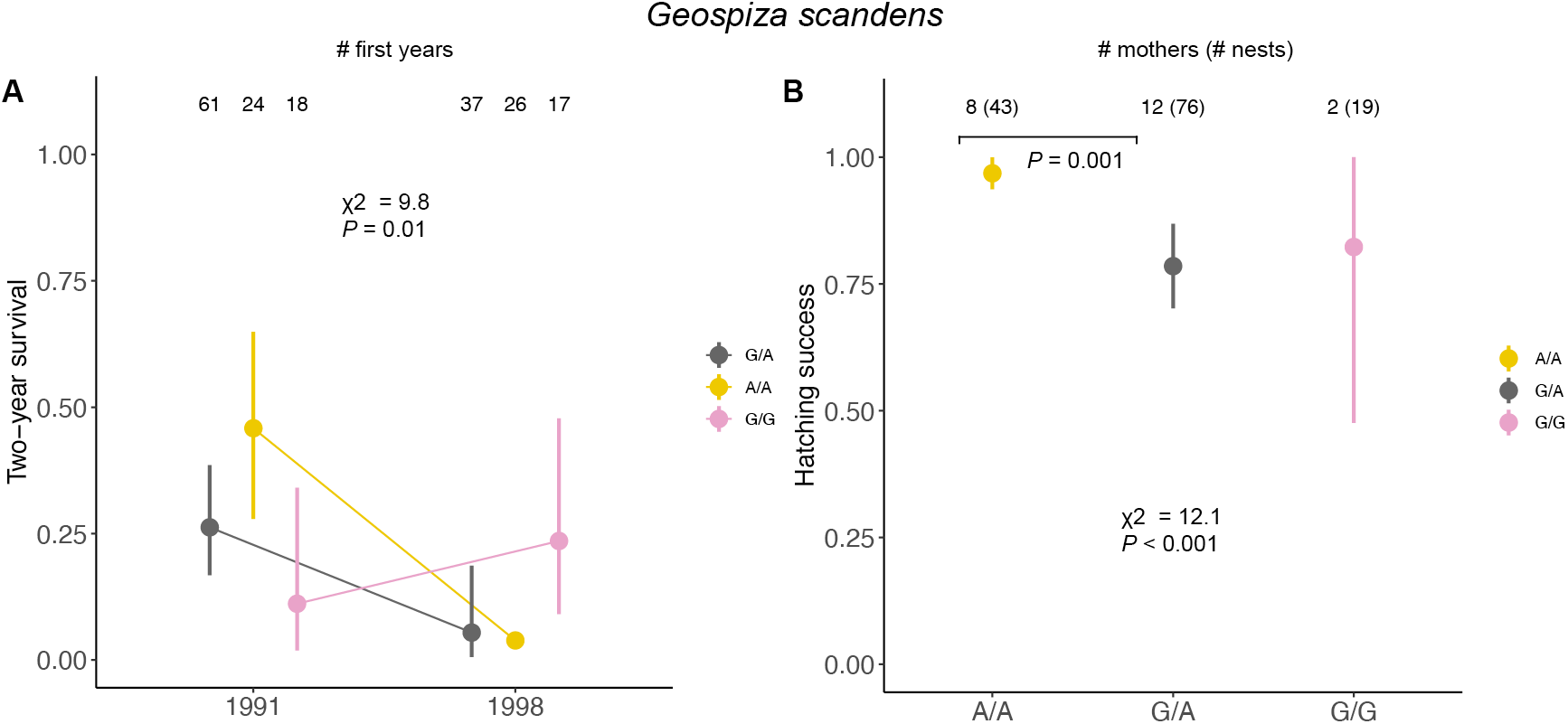
Survival and hatching success in relation to genotype in *G. scandens,* the common cactus finch. **(A)** Differences in survival to 2 years of age between the 1991 cohort and 1998 cohort. Yellow genotype individuals survived better in 1991 than 1998, corresponding to years of high and low cactus availability, respectively. **(B)** Lifetime hatching success for 22 mothers and 138 nests, colored by genotype. Yellow genotype mothers experienced greater hatching success than heterozygotes. Sample sizes are limited as we cannot evaluate homozygous pink hatching success, for which we have only 2 mothers. 95% confidence intervals are shown.

An advantage to yellow genotypes may also extend into adulthood. One possibility is that the polymorphism could affect maternal investment. This is suggested by the finding that chicken mothers homozygous for the yellow leg allele, a *BCO2* allele silenced in skin tissue, are able to invest more carotenoids in egg yolk than other genotypes^31^. In *G. scandens,* yellow genotype mothers hatched eggs more successfully (97%) than heterozygous individuals (78%) when comparing hatching success among 138 nests (χ^2^ = 12.1, *P* < 0.001, df = 2; Fig. 4B). The pattern is repeated at different times and in different age groups in the extended breeding season of 1983 (Extended Data Table 3).

The genetic basis of shared polymorphisms, and the selection pressures acting on them, are generally not known^2^. Here we have shown that variation at a single locus is responsible for a polymorphic color trait in Darwin’s finches. The genetic simplicity of the *BCO2* gene has enabled us to trace the origin of the polymorphism through phylogenetic analysis to a single unique mutation occurring in the Galápagos archipelago approximately half a million years ago.

The polymorphism has been retained in all descendant species except one. The most parsimonious explanation for retention is that the polymorphism is a shared ancestral condition. Persistence across the radiation is remarkable because the rarer morph may be lost through drift in the founding of a new population by a few individuals during speciation^2,6^. Nonetheless, Darwin’s finches satisfy two conditions that are conducive to retention: high speciation rate^20^ and presence of several coexisting, and occasionally interbreeding, closely related species^3,11,32^. Loss through drift is likely to be counteracted by reintroduction through introgression^14,33^, especially in the early stages of speciation. Although we do not fully understand the salient selection pressures, we have identified diet as an important factor. The yellow genotype is relatively common in cactus-specialist species feeding on carotenoid-rich pollen, and they gain a survival advantage in some years when cactus products are plentiful. We found evidence of higher hatching success of eggs produced by females with the homozygous yellow genotype, consistent with a hypothesis that such females deposit more carotenoids in egg yolks than other genotypes.

Most color polymorphisms that have been studied to date are visible in adults and have signaling functions in contexts of mate choice^34^, social dominance^35^, camouflage^7^ or protective mimicry^36^. The yellow beak polymorphism in Darwin’s finches differs from all of these, and is more akin to polymorphisms in Major Histocompatibility Complex (MHC) antigens^33^, where fitness advantages are physiological and biochemical. Since they are largely or entirely out of sight in adults, literally, such polymorphisms may be far more common than is currently recognized, and contribute to diversification in many ways that are yet to be discovered.

## Methods

### Sample collection

Blood was collected as part of a long-term monitoring of finches on Daphne Major and other islands beginning with samples first collected in 1988. Sampling was conducted in accordance with protocols of Princeton University’s Animal Welfare Committee, and stored on EDTA-soaked filter paper in Drierite. Additional details on sample collection can be found elsewhere^3^. We selected 213 individuals for sequencing for which the yellow beak phenotype was known for *G. scandens, G. fortis*, *G. fuliginosa* and hybrids on Daphne Major. We sequenced an equal number (n = 213) of individuals carrying the pink phenotype from the same species on Daphne Major. Embryos (n = 35) were collected on Santa Cruz and Pinta according to^37^. Tissues were stored in RNAlater (ThermoFisher, CA) until further use.

### DNA extraction library preparation

We extracted DNA from blood on filter paper using a custom salt preparation protocol. Briefly, we submerged clipped blood on filter paper in a buffer containing 400mM NaCL, 2mM EDTA (pH 8.0), 10mM TrisHCL (pH 8.0), and dH20. Next, we added a freshly prepared buffer containing 5% SDS, proteinase K (2mg/mL), and dH20. Samples were incubated overnight at 55°C, the filter paper removed, and 135uL of saturated NaCL was added to the mixture. The sample tube was vortexed and spun down at 4,000 rpm for 15min at 4°C and the supernatant transferred to a new 2mL tube. DNA was precipitated using 2 volumes 95% EtOH and mixed by inverting the tube. Finally, samples were spun at 13.3rpm for 3min to pellet the DNA, EtOH removed, and 50-200uL TE added to elute the DNA. DNA concentration was measured on a Nanodrop (ThermoFisher, CA).

We generated fragment libraries for whole-genome sequencing using a custom Tn5-based tagmentation protocol based on^38^. Briefly, we assembled the Tn5 transposon construct using the stock Tn5 (prepared by Karolinska Institutet Protein Science Facility) and primers described in^38^:

Tn5MErev, 59-[phos]CTG TCTCTTATACACATCT-39

Tn5ME-A (Illumina FC-121-1030), 59-TCGTCGGCAGCGTCAGATGTGTATAAGAGACAG-39

Tn5ME-B (Illumina FC-121-1031), 59-GTCTCGTGGGCTCGGAGATGTGTA TAAGAGACAG-39

Samples were tagmented by adding to a solution containing the Tn5 construct and H20, 5x TAPs, and 40% PEG. The mixture was incubated on a Bio-Rad thermocycler (Bio-Rad, CA) for 10min at 55°. Next, genomic libraries were PCR enriched using Kapa Biosystems HiFi HotStart (Wilmington, MA) PCR kit (annealing temperature 63°). DNA libraries were size selected using .38X and .16X Ampure XP beads (Brea, CA) for a target insert size of 350bp, and resulting product quantified on a Tecan microplate reader (Tecan Life Sciences, Switzerland) using Qubit (ThermoFisher, CA) reagents. Samples were pooled in equimolar concentrations and a final size selection performed using .45X and .3X Ampure beads on the resulting pool. Pools were sequenced on an Illumina NovaSeq S4 flow cell (Illumina, CA) with a target sequencing depth of 2x.

### Population genomics

All Illumina short reads were mapped to the chromosome-scale Camarhynchus_parvulus_V1.1 genome assembly (GCA_902806625.1) using BWA mem v0.7.17^39^ and the resulting BAMs were sorted using SAMTOOLS v1.10 (http://www.htslib.org/). Sequencing coverage was estimated for chromosome 4 (a large, representative subset) using the SAMTOOLS coverage command (Extended Data Fig. 1). We used ANGSD v0.931^12^ to estimate genotype likelihoods used for estimating F_ST_. We ran the following commands for each phenotype and chromosome separately:

~~~
$ANGSD_PATH/angsd -b $BAMLIST1 -ref $REFGENOME -anc $REFGENOME -r $INTERVAL \
             -out ${phenotype}_${INTERVAL}.ref \
             -uniqueOnly 1 -remove_bads 1 -only_proper_pairs 0 -trim 0 \
             -minMapQ 20 -minQ 20 -doCounts 1 \
             -GL 1 -doSaf 1 -P 20
~~~

In this step we generated sample allele frequencies per chromosome and all subsequent analyses were run per-chromosome using the chromosome specified by $INTERVAL. We next estimated the 2d folded site frequency spectrum (SFS) using the following command by specifying the

~~~
$ANGSD_PATH/misc/realSFS -P 20 pink_${INTERVAL}.ref.saf.idx
yellow_${INTERVAL}.ref.saf.idx -fold 1 > pink_yellow_${INTERVAL} fold.sfs
~~~

The resulting 2d SFS was used as a prior for estimating F_ST_ first persite then in windows of 10kb (non-overlapping windows). This window-based analysis was carried out for n = 97 pink and n = 97 yellow *G. fortis* samples. These estimates were produced using the following commands:

~~~
$ANGSD_PATH/misc/realSFS fst index_pink_${INTERVAL}.ref.saf.idx
yellow_${INTERVAL}.ref.saf.idx -sfs pink_yellow_${INTERVAL}_fold.sfs -fold 1
-fstout pink_yellow_${INTERVAL}_fold
~~~

~~~
$ANGSD_PATH/misc/realSFS fst stats2 pink_yellow_${INTERVAL}_fold.fst.idx -win
10000 -step 10000 -whichFST 0 > pink_yellow__${INTERVAL}_fold_10kb-
Win.fst.txt
~~~

After the main region of differentiation relating to beak phenotype had been identified, we carried out the above commands for chromosome 24 for all 416 individuals (n = 213 of each phenotype) and the following command to extract per-site F_ST_ estimates.

~~~
$ANGSD_PATH/misc/realSFS fst print
Results_array_SUBSET/${POP1}_${POP2}_${INTERVAL}_fold.fst.idx -r
chr24:4000000-7695000 > ${POP1}_${POP2}_${INTERVAL}_4mb_7.7mb.fst
~~~

All subsequent analysis of F_ST_ estimates were performed using custom scripts in R v3.6.1^40^.

### Analysis of high coverage data

Short read data was accessed from NCBI sequence read archive (www.ncbi.nlm.nih.gov/sra) PRJNA417530, PRJNA263122 and PRJNA301892. Two homozygous pink and homozygous yellow individuals that were sequenced at low coverage were also sequenced to a target coverage of 15x (PRJNA678752). All short-reads were aligned to Camarhynchus_parvulus_V1.1. SNPs were called using GATK’s HaplotypeCaller and joint genotyping using GenotypeGCVFs (v4.1.4.1). Filtering was done for SNPs using filter-expressions in VariantFiltration and only biallelic SNPs were retained:

~~~
“QUAL < 100 | | MQ < 40.0 | | MQ > 80.0 | | MQRankSum < -4.0 | | MQRankSum > 4.0
| | ReadPosRankSum < -4.0 || ReadPosRankSum > 4.0 || QD < 5.0 || FS > 30.0 ||
DP < 50 || DP > 29300”
~~~

And removing genotypes with low depth and low genotype quality using -G-filter:

~~~
“DP < 1 || DP > 200 || GQ < 10”
~~~

### TaqMan Genotyping Assay

Custom SNP genotyping TaqMan assay (ThermoFisher, CA) were applied to perform genotypic analysis of the SNP of interest in exon 4 of *BCO2.* We designed primers (BCO2_F: 5’-TGTTTCAGAACCCAGTGACAACT-3’; BCO2_R: 5’ -TTCCAGTGTCTCTGGGTCCA-3’) and probes (BCO2_VIC:5’-ATGTGAACTACGTGCTGTAC-3’; BCO2_FAM:5’-ATGTGAACTACGTACTGTAC-3’) for the SNP of interest on chr24:6166878 (A/G). We used this assay to genotype 2,890 individuals, for which 1,631 had a known nestling beak phenotype.

### Allele specific expression

We dissected the upper beak primordia of 35 Darwin’s finch embryos (6 *Geospiza magnirostris,* 7 *G. fortis*, 7 *G. fuliginosa*, 8 *Camarhynchus psittacula*, 1 *C. parvulus*, and 6 *Platyspiza crassirostris)* and extracted RNA with E.Z.N.A. Total RNA Kit I (Omega Bio-tek, GA). We prepared cDNA libraries with the NEBNext Ultra RNA Library Prep Kit for Illumina (NewEngland Biolabs, MA) with poly(A) selection. The libraries were then sequenced on HiSeq 4000 (Illumina, CA). For 29 individuals for which we have RNAseq data, we also had genomic DNA available. For these individuals we used the same TaqMan assay to determine genotype. For all other individuals we inferred genotype based on RNA sequencing depth. This includes 1 AA, 4 GG, and 1 GA individuals. Before mapping RNAseq data, Illumina adaptor and primer sequences were removed with CutAdapt v.1.9^41^ and low-quality bases (PHRED < 20) were removed using Trim Galore (v.0.4.1, available at http://www.bioinformatics.babraham.ac.uk/projects/trim_galore/) with default settings. Cleaned RNAseq reads were then mapped to the genome and Fragments Per Kilobase of transcript per Million mapped reads (FPKM) was calculated using the HISAT2 (v. 2.1.0) - StringTie (v. 1.3.6) - Ballgown (v. 2.20.0) pipeline^42^. For the 6 individuals we genotyped using RNAseq data, genotypes were called using SAMTOOLS and BCFtools v1. 9:

~~~
samtools mpileup -Q 20 -q 20 -t DP4,DP -vuf ${ref} *.bam | bcftools call -M -
f GQ -mg 3 -Ov > snps.vcf
~~~

We compared the number of Fragments Per Kilobase of transcript per Million mapped reads using a three-way ANOVA with the aov command and post-hoc comparisons using the TukeyHSD command in R (v3.6.1).

In order to search for splice variations, RT-PCR was applied to amplify the regions around the SNP in cDNA by the use of forward 5’-CCCATCCCAGCCAAGATCAA-3’ and reverse 5’-CGTAGTGGGGATGAGCTGTG-3’ primers under the following conditions: 95 °C for 5 min, followed by 35 cycles of 95°C for 30 s, 60°C for 30 s and 72°C for 1 min. The amplified fragments were subjected to Sanger sequencing.

### Survival analysis and hatching success

We modeled early life survival using a generalized linear model (GLM) in R (v3.6.1) with survival to their second year (Age = 2) as a binary response variable. We analyzed 2-year survival to capture the consequences of back-to-back cactus fruit reduction beginning in 1998. We analyzed early year survival in 1991 and 1998 using a GLM with the *BCO2* genotype, year, and the interaction between the two to test the hypothesis that first year survival differed between 1991 and 1998. We used the Anova command (type = “II”) from the car package^43^ to evaluate significance of predictor effects.

In order to evaluate the immune defense hypothesis, we ran a logistic regression using GLM in R (v3.6.1) on “survived” or “died” (survival from 2008 to 2009) and the individual’s BCO2 genotype, pox infection (YES or NO) and age. Pox infection status for the year 2008 was taken from^28^. We additionally included the interaction between BCO2 genotype and Pox to test for a differential effect of genotype on immune response to pox. We did not find an interaction between pox infection status and *BCO2* genotype when evaluating the effect of predictors using the car packageAnova (type = “II”) command, but yellow homozygotes survived poorly in 2008 irrespective of infection status.

We analyzed lifetime mean female hatching success in *G. scandens* using linear mixed models in the package lmerTest ^44^ Mean hatching was calculated per nest as the number of eggs that hatched and was only tabulated for individuals where the number of eggs laid and hatched were known. We removed nests where no eggs hatched (which could have been the result of other factors, such as nest predation) and birds with only one nest (which prevented a reliably mean rate across nests). One yellow phenotype individual was included as AA who failed to amplify using the TaqMan assay. Each individual was given a single value for lifetime average hatching success, which was used as the response variable in the LMM. We included cohort (year born) of each female as a random effect in the model. We used the Anova command (type = “II”) from the car package ^43^ to test significance of predictor effects.

Additional survival analyses, hatching success, and fledging success can be found in the supplemental R markdown notebook “BCO2 stats” at https://github.com/erikenbody/Finch_beak_color_polymorphism.

## Acknowledgments

The National Genomics Infrastructure (NGI)/Uppsala Genome Center and UPPMAX provided service in massive parallel sequencing and computational infrastructure. A. Sendell-Price provided comments on an earlier draft. **Funding:** NSF (USA) funded the collection of material under permits from the Galápagos and Costa Rica National Parks Services and the Charles Darwin Research Station, and in accordance with protocols of Princeton University’s Animal Welfare Committee. The study was supported by The Knut and Alice Wallenberg Foundation and Vetenskapsrådet.

## Author contributions

LA, PRG, BRG, and EDE conceived and designed the study. PRG and BRG collected blood samples and field observations. EDE conducted all bioinformatic and ecological analyses with input from PRG and BRG. CGS was responsible for all genomic material preparation, sequencing, and pedigree analysis. CGS and HB conducted genotyping. MD, OO, and AA collected samples and generated expression data. CJR generated the genome assembly. EDE wrote the first draft of the manuscript with input from LA, PRG, and BRG and all authors commented on the final manuscript. All authors approved the manuscript before submission.

## Data availability

All novel sequencing data from this study will be made available through NCBI Sequence Read Archive BioProject ID PRJNA678752.

## Code availability

All code used to analyze sequence data will be uploaded to the Github page of EDE before publication (https://github.com/erikenbody/Finch_beak_color_polymorphism).

## Materials and correspondence

Erik D. Enbody and Leif Andersson

## Extended data

### Supplementary text

#### Notes on hybridization between *G. fortis* and *G. scandens*

Introgressive hybridization of *G. fortis* and *G. scandens* carries little or no fitness penalty and genes are exchanged bidirectionally but unequally: *G. fortis* donates more often than it receives^22^. The frequency of hybrids (F_1_ and backcrosses), as recognized by microsatellite assignments^22,45^, varied in the *G. fortis* population from 0.03 to 0.15, and in the *G. scandens* population from 0.03 to 0.32.

In *G. fortis*, As with *G. scandens* (main text), frequencies of homozygotes *BCO2* covary negatively (adj R^2^ = 0.25, *P* = 0.006). The frequency of the common genotype (homozygous pink) negatively covaries with the frequency of hybrids (adj R^2^ = 0.30, *P* = 0.0025), whereas the frequency of the rarer yellow genotype does not (adj R^2^ = 0.03, *P*>0.1).

In *G. fortis,* coefficients of annual variation are 17.3 ± 5.0: 95 percent confidence limits) for heterozygotes and 20.0 ± 5.8 for AA genotypes and 29.4 ± 8.8 for GG homozygotes. Annual variation in the frequencies of heterozygotes is positively related to the frequencies of hybrids in the sample (adj R^2^ = 0.26, P = 0.0048). Frequencies of GG homozygotes vary negatively with hybrid frequency (adj R^2^ = 0.30, P = 0.0025), which is expected because the G allele has a lower frequency in the donor species, *G. scandens.* Frequencies of AA genotypes are low and vary independently of hybrid frequencies (adj R^2^ = 0.03, P>0.1). *G. fortis* (but not *G. scandens)* occasionally hybridizes with immigrant *G. fuliginosa*. The hybrids they produce have no effect on the above results when included in the calculations.

**Extended Data Fig. 1:**
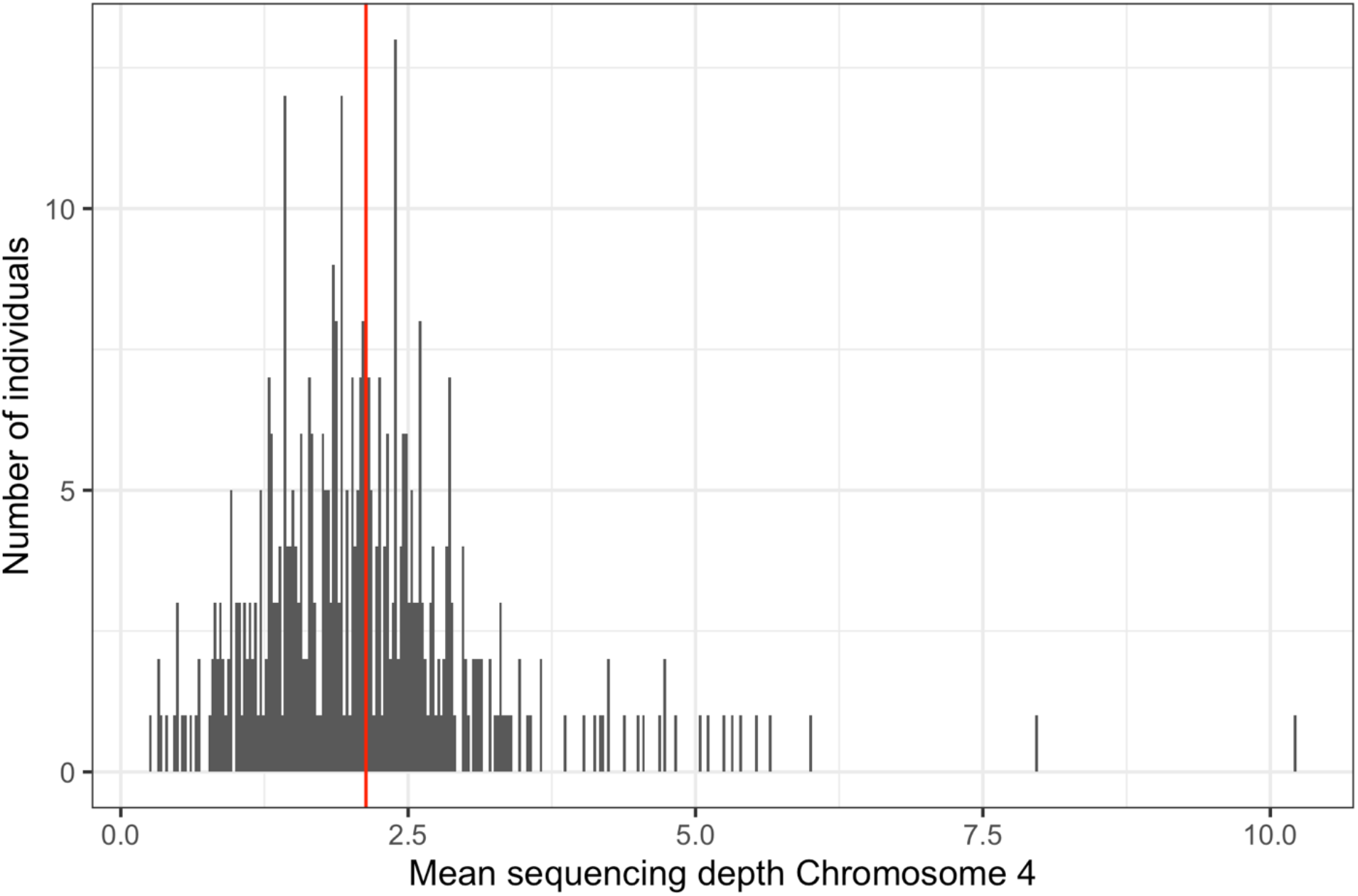
Mean sequencing depth across Chromosome 4 for all 426 samples used in the F_ST_ contrast in Fig. 1B. Average sequencing depth across this chromosome (2.1X) is marked with a red vertical line. Sequencing depth was calculated using the SAMTOOLS v1.10 (http://www.htslib.org/) coverage command.

**Extended Data Fig. 2:**
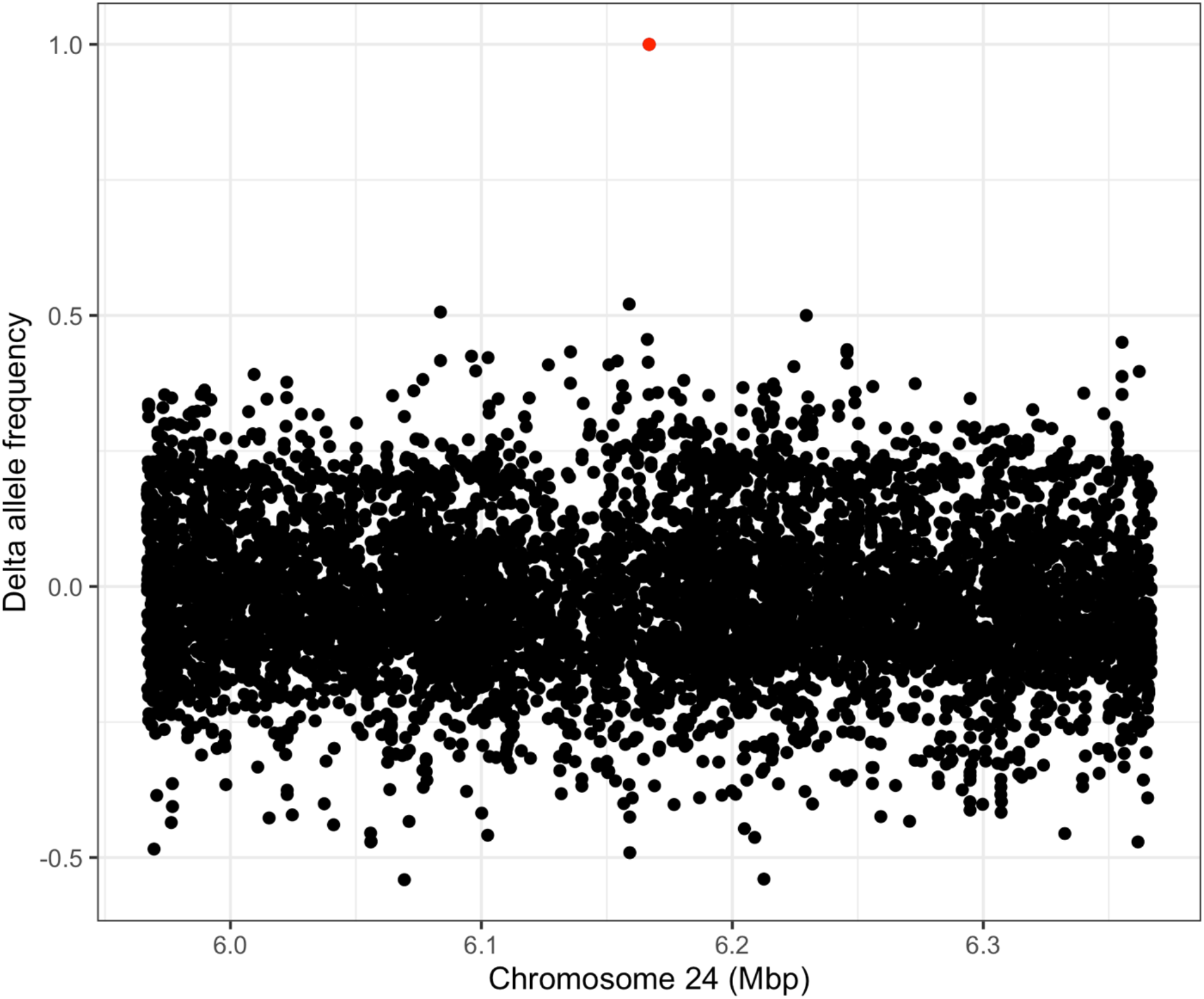
Concise overview of our comprehensive search for linked variants (both SNPs and indels) in high coverage individuals. Here, all SNPs and indels 200kbp upstream and downstream of the SNP of interest (6166878) are included and allele frequency calculated for homozygous alternative individuals at this position. Delta allele frequency is calculated as: (frequency of alternate genotype in homozygous yellow genotype individuals) - (frequency of alternate genotype in homozygous pink genotype individuals). Only the synonymous SNP in *BCO2* exon 4 is consistently different between these group, suggesting it is not in linkage disequilibrium with another causal SNP or indel.

**Extended Data Fig. 3:**
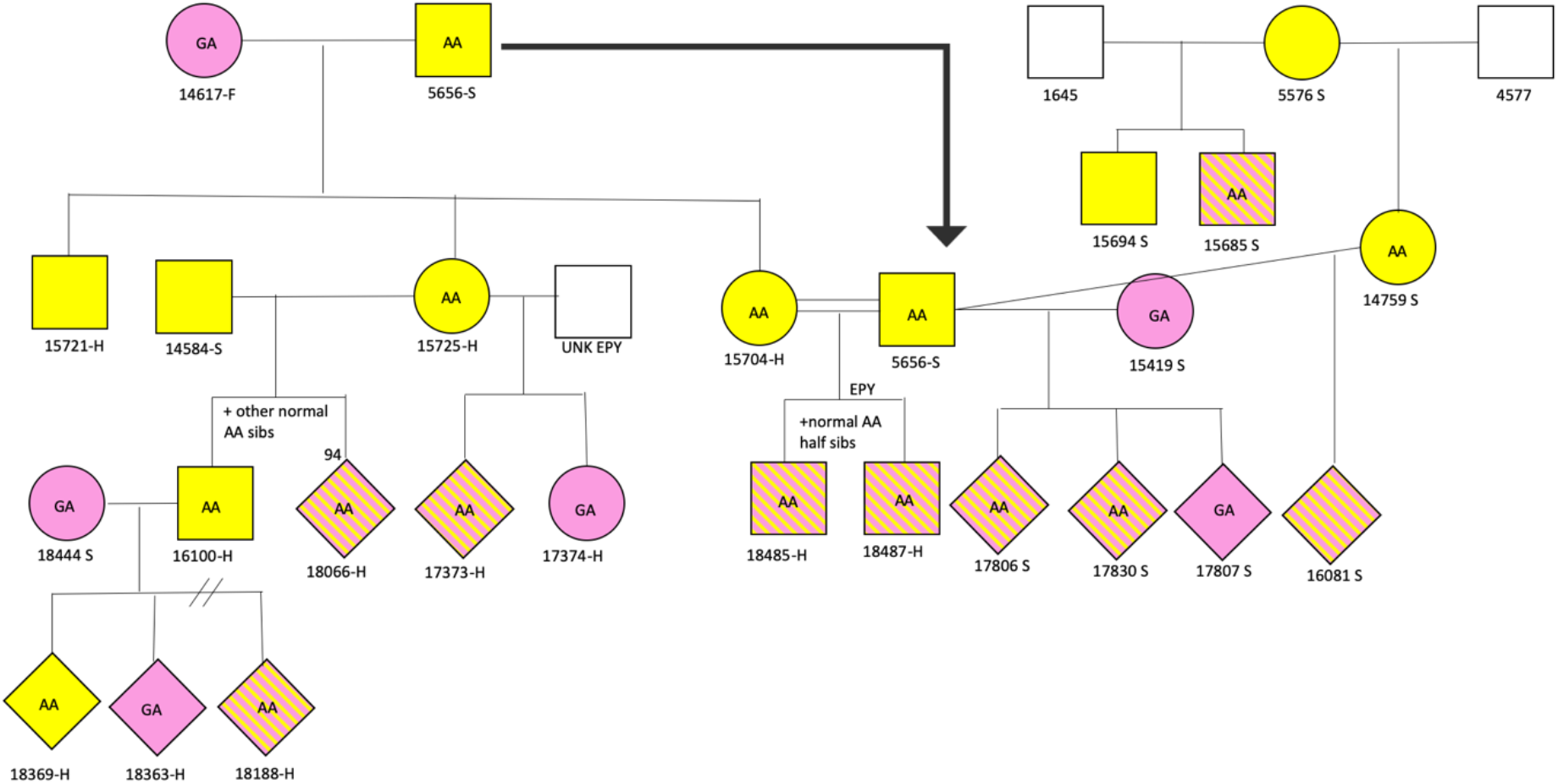
Simplified pedigree of anomalous individuals (conflicting genotype and phenotype). Known genotype individuals are marked as AA or GA and colored by their phenotype yellow or pink. AA individuals with a pink phenotype are marked with hatched lines. Note that not all siblings and mates are shown, but 1/3 of all anomalous AA individuals are represented in this pedigree. The number below each individual is its unique band combination and the suffice refers to species (S = *G. scandens,* F = *G. fortis,* H = hybrid). Squares represent males, circles represent females, and diamonds are unknown sex.

**Extended Data Fig. 4:**
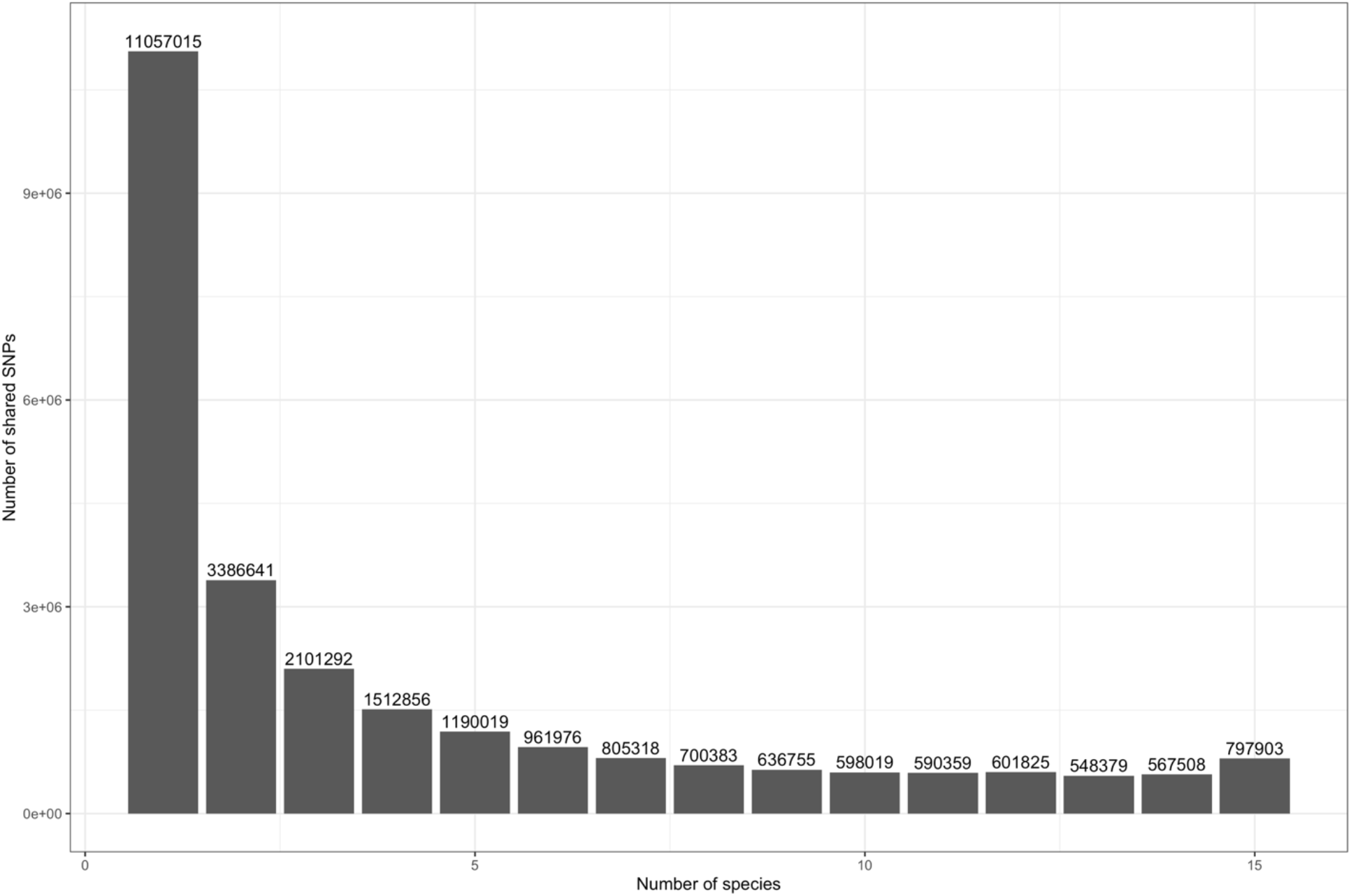
Allele sharing among species of Darwin’s finches. Polymorphic sites that segregate in all ground, tree, and Cocos finch were filtered for missing data (data in 90% of all individuals) and tabulated for all variable sites among the 15 species carrying the yellow (A) allele and *G. septentrionalis.* The number of sites shared between x number of species (x-axis) were tabulated and plotted as a histogram here. Approximately 5% of all polymorphic sites are shared in at least 14 species.

**Extended Data Fig. 5:**
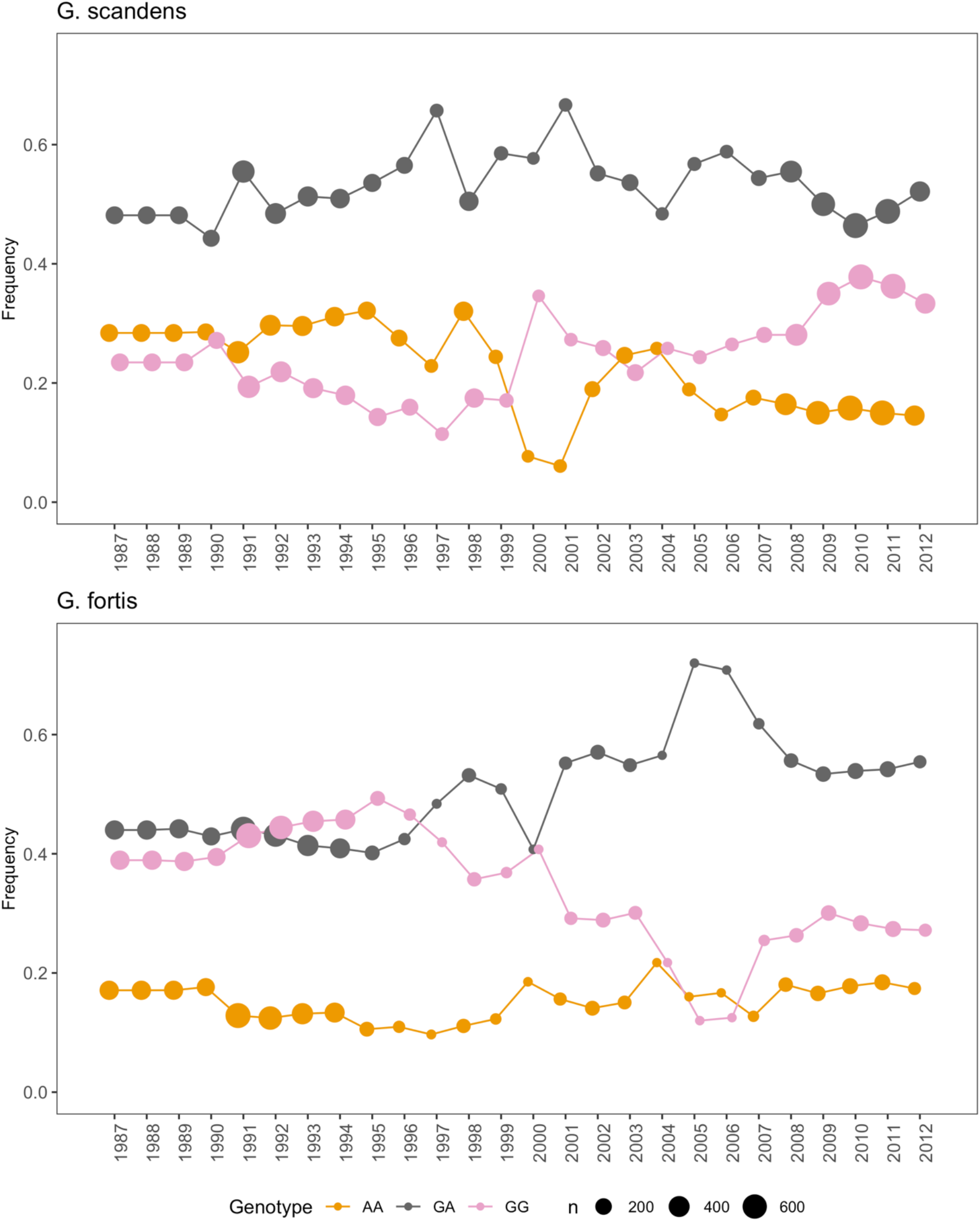
Frequency of pink homozygotes (pink), yellow homozygotes (yellow), and heterozygotes (grey) across years for *G. scandens* (top) and *G. fortis* (bottom). The size of each point reflects the number of adults alive that year.

**Extended Data Fig. 6:**
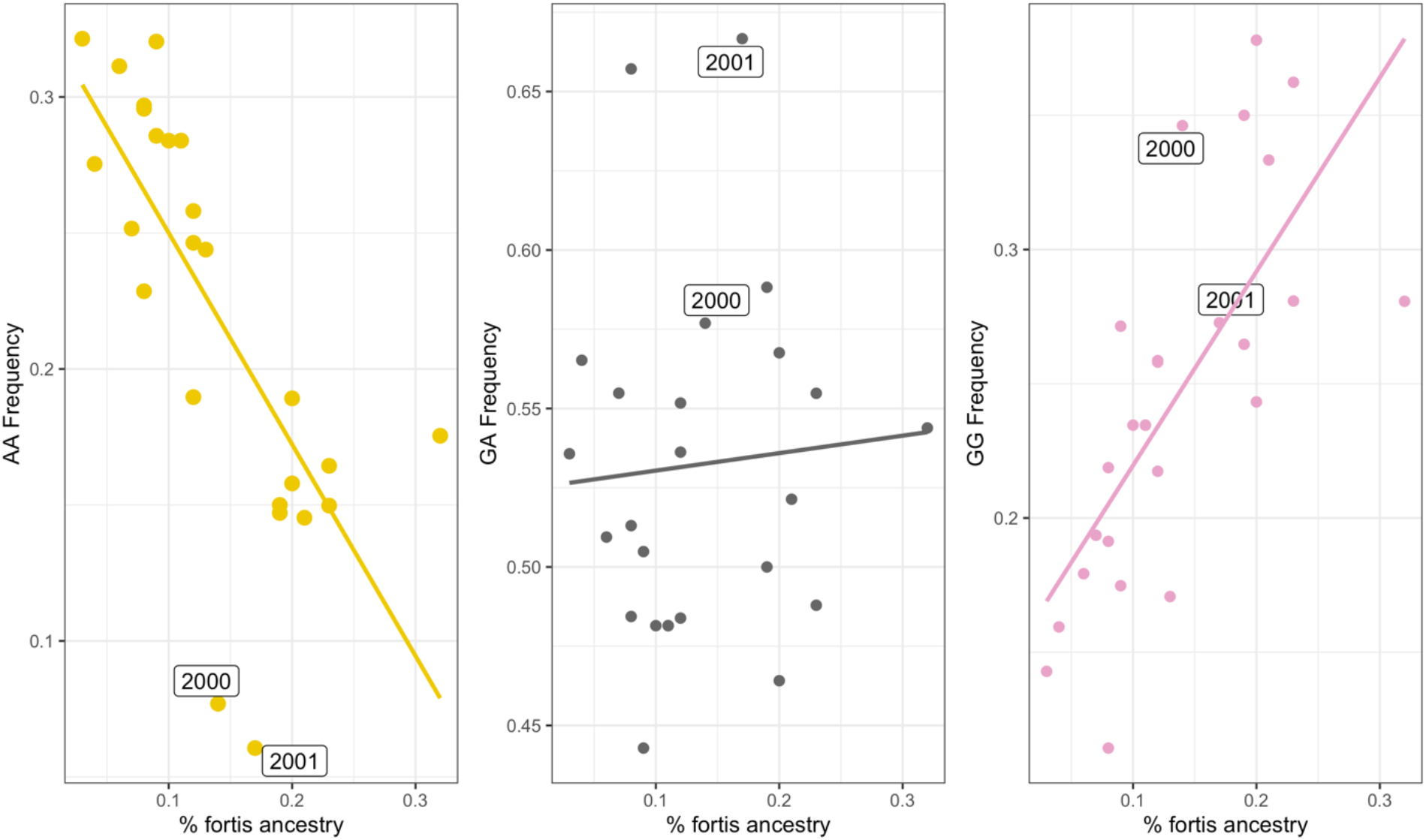
Percent *G. fortis* ancestry in *G. scandens.* Each dot represents one year. R^2^ coefficients are as follows: AA = −0.71, GA = 0.07, GG = 0.72.

**Extended Data Fig. 7:**
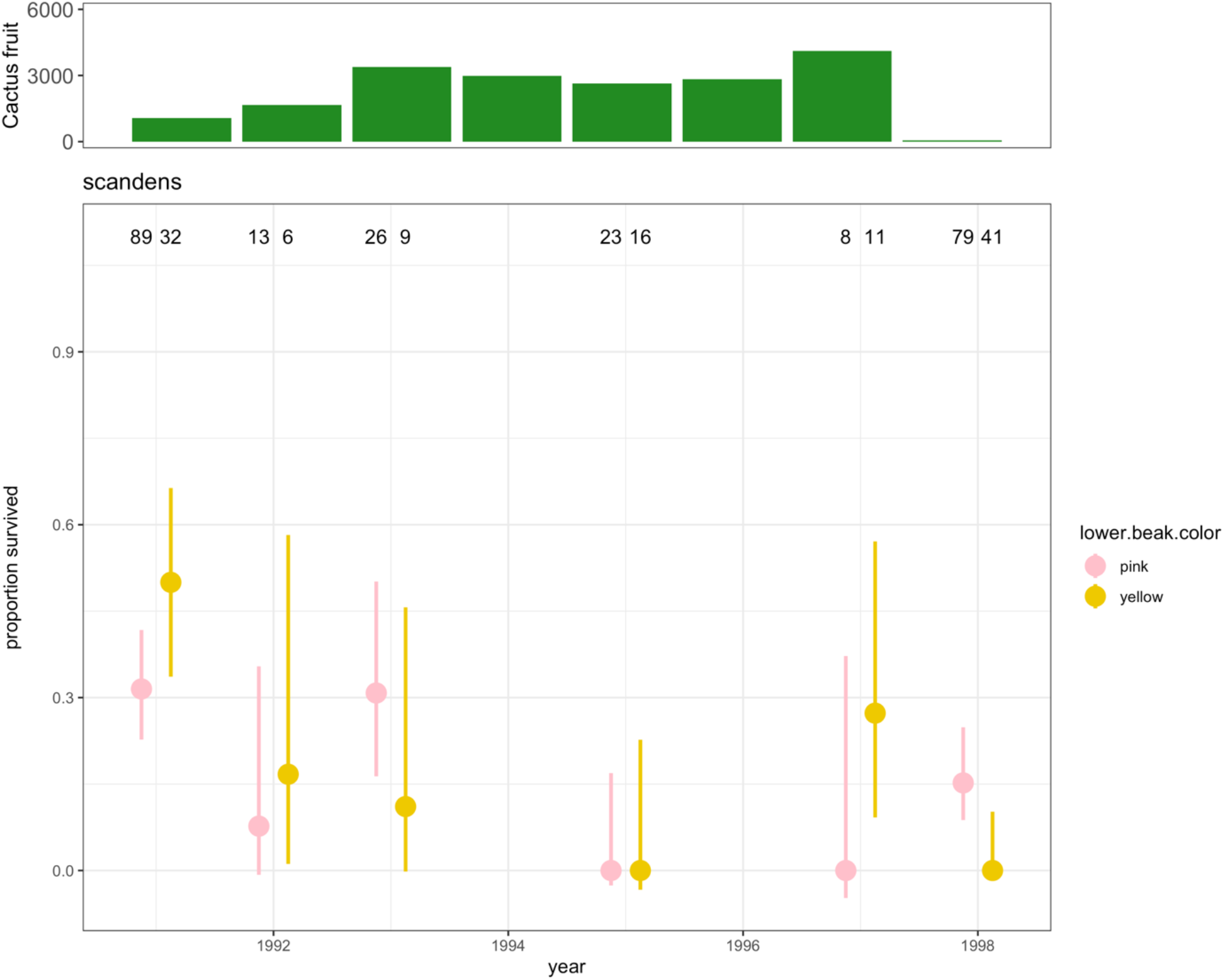
*G. scandens* survival to second year after fledgling, grouped by nestling phenotype. Between 1991 and 1998 most years the yellow phenotype survived better. The difference between years 1991 and 1998 match differences in survival reported using genotype data. 95% confidence intervals are shown.

**Extended Data Fig. 8:**
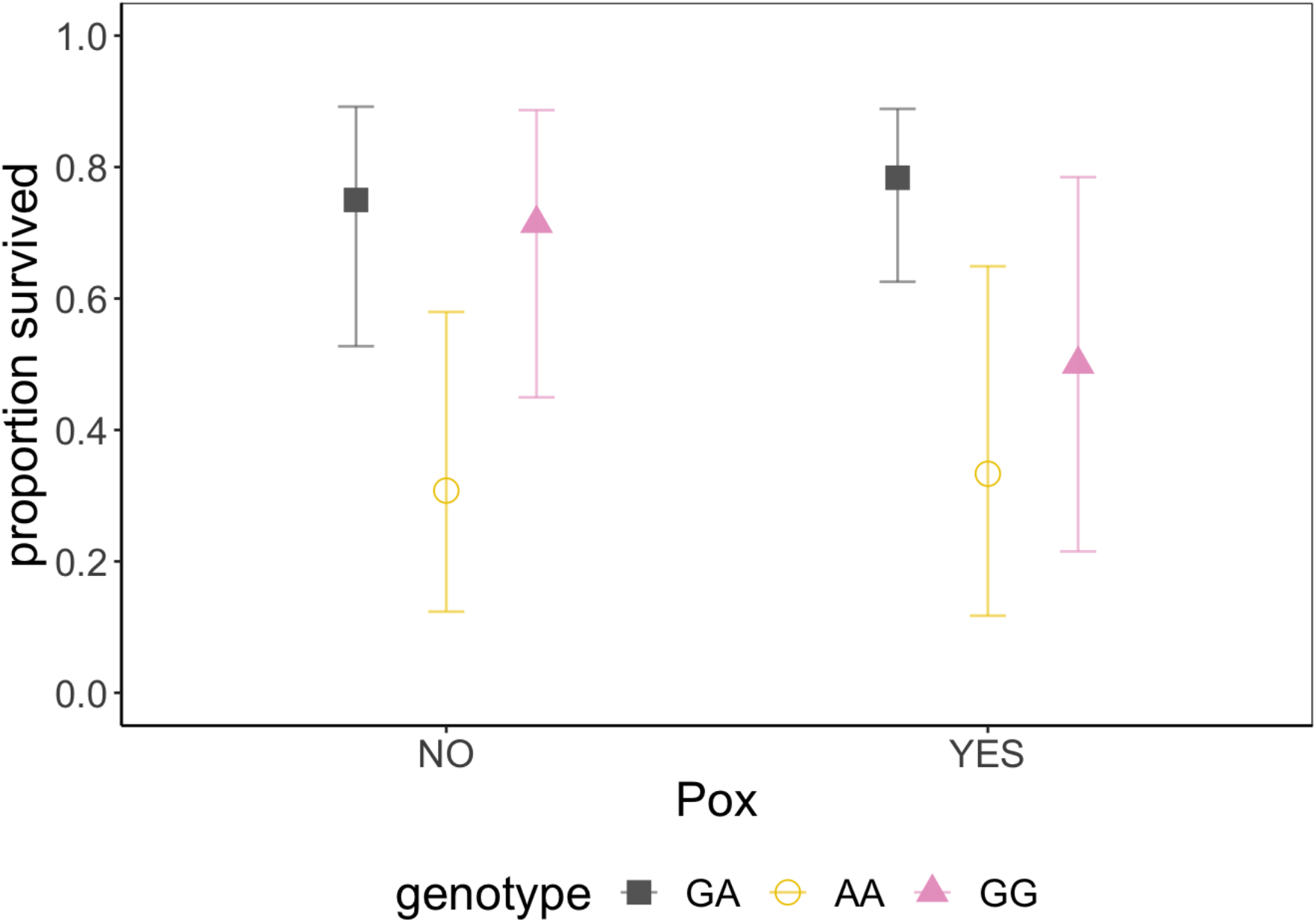
The proportion of pox infected birds that survived to the following year, separated by *BCO2* genotype. The sample sizes for pox infected individuals were (AA=9, GA=37, GG=8) and for non-infected individuals (AA=13, GA=20, GG=14). We found no interaction between pox infection and *BCO2* genotype (χ^2^ =0.98, *P* = 0.62, Materials and Methods). 95% confidence intervals are shown.

**Extended Data Fig. 9:**
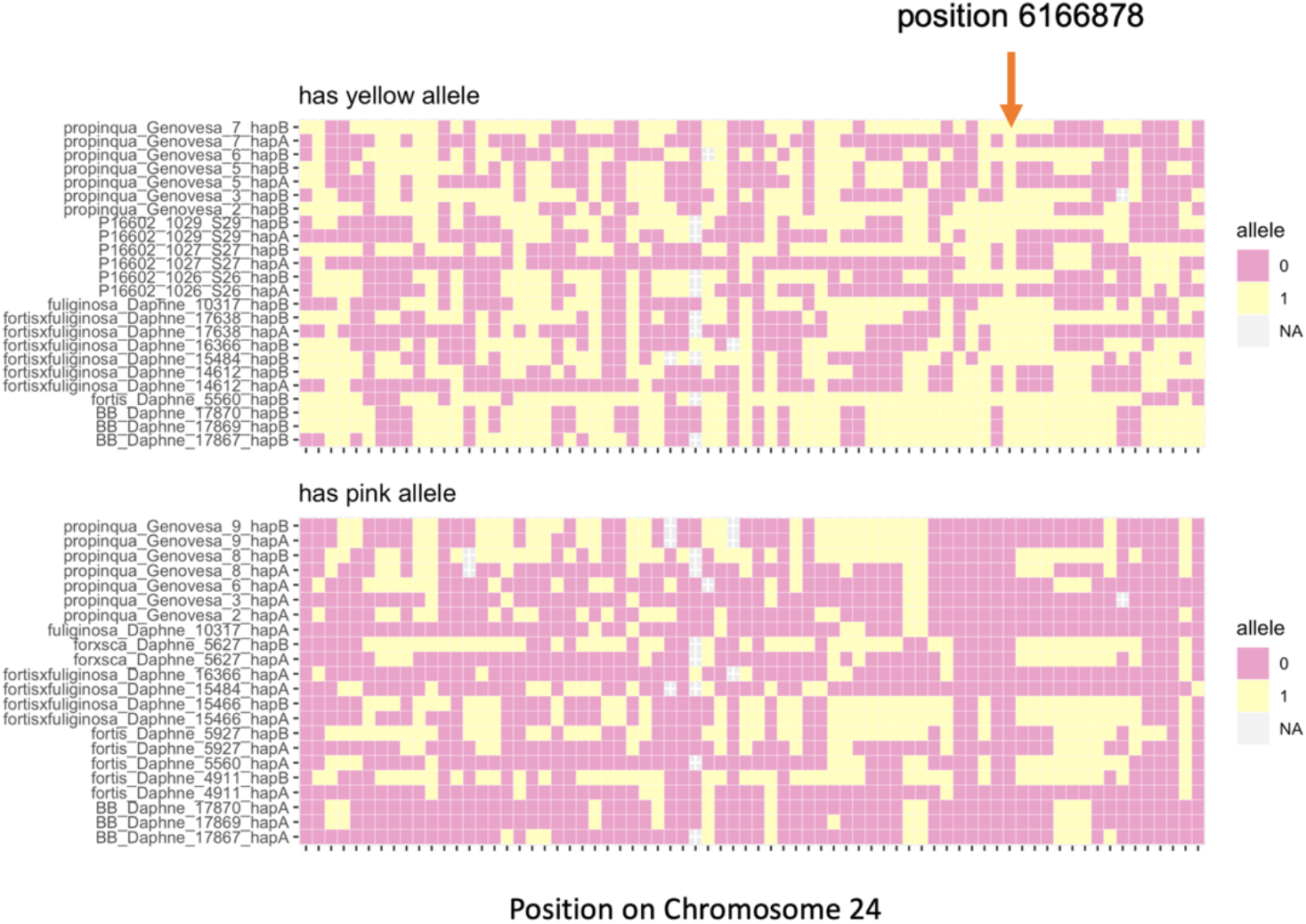
Haplotype blocks for high coverage sequenced individuals carrying the pink or yellow allele. Only high F_ST_ SNPs are shown (between yellow and pink phenotypes) across a 440kbp region. Only the SNP of interest shows a perfect association with the two alleles (marked with a red arrow above). Some haplotype structure is present in the vicinity of *BCO2,* but no other variants show strong association with the identified SNP of interest. Together this suggests that the yellow allele occurs on multiple haplotypes across different species.

**Extended Data Table 1:** Summary of phenotypes for 1,631 individuals checked using the TaqMan assay for the *BCO2* SNP identified in this study.

| **Species** | **Island** | **Beak Color** | **n** |
| --- | --- | --- | --- |
| propinqua | Genovesa | Pink | 5 |
| propinqua | Genovesa | Yellow | 1 |
| fortis | Daphne | Pink | 774 |
| fortis | Daphne | Yellow | 112 |
| fuliginosa | Daphne | Pink | 5 |
| hybrid | Daphne | Pink | 364 |
| hybrid | Daphne | Yellow | 104 |
| magnirostris | Daphne | Pink | 1 |
| scandens | Daphne | Pink | 196 |
| scandens | Daphne | Yellow | 69 |

**Extended Data Table 2:**
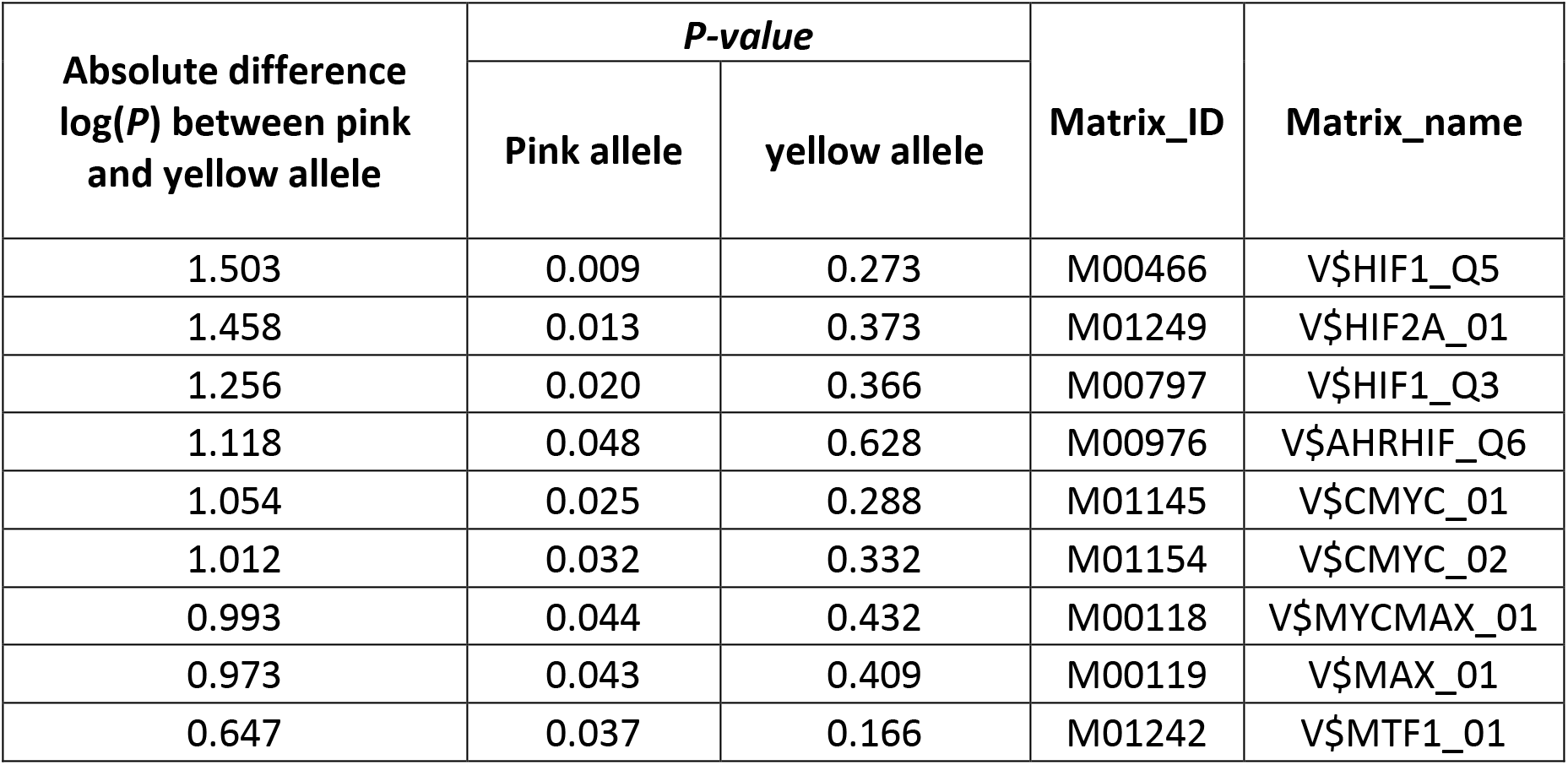
sTRAP^19^ analysis for predicted differential transcription factor affinities involving the SNP of interest in *BCO2* exon 4. *P*-values for the pink and yellow allele are shown, as well as the absolute different in in transcription factor binding probability for the two alleles. The input for sTRAP was 100bp of sequence flanking the SNP of interest and the only difference between the two sequences being the SNP of interest.

| Absolute difference<br>log(*P*) between pink<br>and yellow allele | *P-value* |  | Matrix_ID | Matrix_name |
| --- | --- | --- | --- | --- |
|  | Pink allele | yellow allele |  |  |
| 1.503 | 0.009 | 0.273 | M00466 | V\$HIF1_Q5 |
| 1.458 | 0.013 | 0.373 | M01249 | V\$HIF2A_01 |
| 1.256 | 0.020 | 0.366 | M00797 | V\$HIF1_Q3 |
| 1.118 | 0.048 | 0.628 | M00976 | V\$AHRHIF_Q6 |
| 1.054 | 0.025 | 0.288 | M01145 | V\$CMYC_01 |
| 1.012 | 0.032 | 0.332 | M01154 | V\$CMYC_02 |
| 0.993 | 0.044 | 0.432 | M00118 | V\$MYCMAX_01 |
| 0.973 | 0.043 | 0.409 | M00119 | V\$MAX_01 |
| 0.647 | 0.037 | 0.166 | M01242 | V\$MTF1_01 |

**Extended Data Table 3:** Hatching success in 1983. An extended breeding season in 1983 allowed us to compare hatching success variation between different phenotypes within a single season under different ecological conditions. Breeding success was uniformly high in the first 3 months of 1983, and then low following extensive rains and the Early period refers to the first three months. Cactus flowers were present in the first half, dwindling to zero, and absent in the second half. Yellow phenotype individuals experienced greater hatching success regardless of conditions and mirror the results shown for lifetime hatching success in the main text using a much larger sample size with genotype data. Red text marks the higher hatching success per species.

| Species | Phenotype | Hatch success | SD | N mothers |
| --- | --- | --- | --- | --- |
| EARLY FIRST HALF |  |  |  |  |
| *fortis* | pink | 0.76 | 0.20 | 33 |
| *fortis* | yellow | 0.83 | 0.14 | 9 |
| *scandens* | pink | 0.71 | 0.18 | 11 |
| *scandens* | yellow | 0.86 | 0.20 | 5 |
| EARLY SECOND HALF |  |  |  |  |
| *fortis* | pink | 0.74 | 0.18 | 32 |
| *fortis* | yellow | 0.88 | 0.15 | 8 |
| *scandens* | pink | 0.69 | 0.23 | 10 |
| *scandens* | yellow | 0.77 | 0.15 | 5 |

## References

1. Gray, S. M. & McKinnon, J. S. Linking color polymorphism maintenance and speciation. Trends Ecol. Evol. 22, 71–79 (2007).

2. Jamie, G. A. & Meier, J. I. The Persistence of Polymorphisms across Species Radiations. Trends Ecol. Evol. 35, 795–808 (2020).

3. Grant, P. R. & Grant, B. R. 40 years of evolution: Darwin’s Finches on Daphne Major Island. 40 Years of Evolution: Darwin’s Finches on Daphne Major Island (Princeton University Press, 2014). doi:10.5860/choice.52-0821.

4. Schluter, D. The Ecology of Adaptive Radiation. (Oxford University Press, 2000).

5. Stroud, J. T. & Losos, J. B. Ecological Opportunity and Adaptive Radiation. Annu. Rev. Ecol. Evol. Syst. 47, 507–532 (2016).

6. Guerrero, R. F. & Hahn, M. W. Speciation as a Sieve for Ancestral Polymorphism. Mol. Ecol. 38, 42–49 (2017).

7. Nosil, P. et al. Natural selection and the predictability of evolution in timema stick insects. Science 359, 765–770 (2018).

8. Llaurens, V., Whibley, A. & Joron, M. Genetic architecture and balancing selection: the life and death of differentiated variants. Mol. Ecol. 26, 2430–2448 (2017).

9. Villoutreix, R. et al. Large-scale mutation in the evolution of a gene complex for cryptic coloration. Science 369, 460–466 (2020).

10. Grant, P. R., Boag, P. T. & Schluter, D. A Bill Color Polymorphism in Young Darwin’s Finches. Auk 96, 800–802 (1979).

11. Grant, B. R. & Grant, P. R. Evolutionary Dynamics of a Natural Population: The Large Cactus Finch of the Galapagos. (University of Chicago Press, 1989).

12. Korneliussen, T. S., Albrechtsen, A. & Nielsen, R. ANGSD: Analysis of Next Generation Sequencing Data. BMC Bioinformatics 15, 356 (2014).

13. Eriksson, J. et al. Identification of the Yellow skin gene reveals a hybrid origin of the domestic chicken. PLoS Genet. 4, (2008).

14. Andrade, P. et al. Regulatory changes in pterin and carotenoid genes underlie balanced color polymorphisms in the wall lizard. Proc. Natl. Acad. Sci. U. S. A. 116, 5633–5642 (2019).

15. Våge, D. I. & Boman, I. A. A nonsense mutation in the beta-carotene oxygenase 2 (BCO2) gene is tightly associated with accumulation of carotenoids in adipose tissue in sheep *(Ovis aries)*. BMC Genet. 11, 10 (2010).

16. Hill, G. E. A Red Bird in a Brown Bag: The Function and Evolution of Colorful Plumage in the House Finch. (Oxford University Press, 2002).

17. Machado, H. E., Lawrie, D. S. & Petrov, D. A. Pervasive Strong Selection at the Level of Codon Usage Bias in Drosophila melanogaster. Genetics 214, 511–528 (2020).

18. Zhou, Z. et al. Codon usage is an important determinant of gene expression levels largely through its effects on transcription. Proc. Natl. Acad. Sci. 113, E6117–E6125 (2016).

19. Thomas-Chollier, M. et al. Transcription factor binding predictions using TRAP for the analysis of ChIP-seq data and regulatory SNPs. Nat. Protoc. 6, 1860–1869 (2011).

20. Lamichhaney, S. et al. Evolution of Darwin’s finches and their beaks revealed by genome sequencing. Nature 518, 371–375 (2015).

21. Lamichhaney, S. et al. Rapid hybrid speciation in Darwin’s finches. Science 359, 224–228 (2018).

22. Grant, P. R. & Grant, B. R. Hybridization increases population variation during adaptive radiation. Proc. Natl. Acad. Sci. U. S. A. 116, 23216–23224 (2019).

23. Grant, B. R. Pollen Digestion by Darwin’s Finches and Its Importance for Early Breeding. Ecology 77, 489–499 (1996).

24. Kiefer, C. et al. Identification and Characterization of a Mammalian Enzyme Catalyzing the Asymmetric Oxidative Cleavage of Provitamin A. J. Biol. Chem. 276, 14110–14116 (2001).

25. Lyon, B. E., Eadie, J. M. & Hamilton, L. D. Parental choice selects for ornamental plumage in American coot chicks. Nature 371, 240–243 (1994).

26. Blount, J. D., Metcalfe, N. B., Birkhead, T. R. & Surai, P. F. Carotenoid modulation of immune function and sexual attractiveness in zebra finches. Science 300, 125–127 (2003).

27. Lobo, G. P., Isken, A., Hoff, S., Babino, D. & von Lintig, J. BCDO2 acts as a carotenoid scavenger and gatekeeper for the mitochondrial apoptotic pathway. Dev. 139, 2966–2977 (2012).

28. Huber, S. K. et al. Ecoimmunity in Darwin’s finches: Invasive parasites trigger acquired immunity in the medium ground finch *(Geospiza fortis)*. PLoS One 5, (2010).

29. McGraw, K. J. & Toomey, M. B. Carotenoid accumulation in the tissues of zebra finches: Predictors of integumentary pigmentation and implications for carotenoid allocation strategies. Physiol. Biochem. Zool. 83, 97–109 (2010).

30. Huggins, K. A., Navara, K. J., Mendonça, M. T. & Hill, G. E. Detrimental effects of carotenoid pigments: the dark side of bright coloration. Naturwissenschaften 97, 637–644 (2010).

31. Fallahshahroudi, A., Sorato, E., Altimiras, J. & Jensen, P. The Domestic BCO2 Allele Buffers LowCarotenoid Diets in Chickens: Possible Fitness Increase Through Species Hybridization. Genetics 212, 1445–1452 (2019).

32. Grant, P. R., Grant, B. R. & Petren, K. Hybridization in the Recent Past. Am. Nat. 166, 56–67 (2005).

33. Hedrick, P. W. Adaptive introgression in animals: Examples and comparison to new mutation and standing variation as sources of adaptive variation. Mol. Ecol. 22, 4606–4618 (2013).

34. Lamichhaney, S. et al. Structural genomic changes underlie alternative reproductive strategies in the ruff *(Philomachuspugnax)*. Nat. Genet. 48, 84–88 (2015).

35. Dey, C. J., Valcu, M., Kempenaers, B. & Dale, J. Carotenoid-based bill coloration functions as a social, not sexual, signal in songbirds (Aves: Passeriformes). J. Evol. Biol. 28, 250–8 (2015).

36. Jay, P. et al. Supergene Evolution Triggered by the Introgression of a Chromosomal Inversion. Curr. Biol. 28, 1839–1845.e3 (2018).

37. Abzhanov, A. Collection of embryos from Darwin’s finches (Thraupidae, Passeriformes). Cold Spring Harb. Protoc. 4, (2009).

38. Picelli, S. et al. Tn5 transposase and tagmentation procedures for massively scaled sequencing projects. Genome Res. 24, 2033–2040 (2014).

39. Li, H. & Durbin, R. Fast and accurate long-read alignment with Burrows-Wheeler transform. Bioinformatics 26, 589–595 (2010).

40. R Core Team. R: A Language and Environment for Statistical Computing. (2019).

41. Martin, M. Cutadapt removes adapter sequences from high-throughput sequencing reads. EMBnet.journal 17, 10 (2011).

42. Pertea, M., Kim, D., Pertea, G. M., Leek, J. T. & Salzberg, S. L. Transcript-level expression analysis of RNA-seq experiments with HISAT, StringTie and Ballgown. Nat. Protoc. 11, 1650–1667 (2016).

43. Fox, J. & Weisberg, S. An {R} Companion to Applied Regression, Second Edition. (SAGE publications, 2011).

44. Kuznetsova, A., Brockhoff, P. B. & Christensen, R. H. B. lmerTest Package: Tests in Linear Mixed Effects Models. J. Stat. Softw. 82, (2017).

45. Grant, P. R. & Grant, B. R. Conspecific versus heterospecific gene exchange between populations of Darwin’s finches. Philos. Trans. R. Soc. B Biol. Sci. 365, 1065–1076 (2010).

